# Development of glowing plastic bricks from waste plastic and florescent pyoverdine pigment of *Pseudomonas aeruginosa*

**DOI:** 10.1101/2024.11.05.622189

**Authors:** Kshipra Pandey, Rushita Oderdara, Ritu Patel

**Affiliations:** Department of Bioscience, Indrashil University, Rajpur, Gujarat, India 382740

**Keywords:** Glowing bricks, *Pseudomonas aeruginosa*, Plastic waste, Pyoveridine Pigment, Recycle

## Abstract

The extensive increase in non-biodegradable plastic waste is leading to heightened pollution and severe impacts on the climate. The marine environment withstands the worst of plastic pollution, with a documented accumulation of 86 million tons of plastic debris globally. Humans are directly impacted by degraded plastic waste through direct consumption (e.g., in tap water), indirect consumption (via the consumption of contaminated animals), and interference with various hormonal mechanisms. In the 20th century, human-produced plastics constituted approximately 3% of the world’s total biomass. In addressing this issue, we embarked on the development of plastic bricks, emphasizing the repurposing of plastic waste. To distinguish our approach, we incorporated a luminescent quality using the inherent fluorescent property of the pyoverdine pigment produced by *Pseudomonas aeruginosa*. This luminescence aims to mitigate nocturnal visibility issues, which contribute to accidents involving both human and speechless fauna.

## Introduction

Plastic pollution is a worldwide issue. Every year 19-23 million tonnes of plastic waste leaks into rivers, polluting lakes, aquatic ecosystems, and seas (https://www.unep.org/plastic-pollution). The world’s oceans, lakes, and rivers are polluted with the equivalent of 2,000 garbage trucks of plastic waste daily. Plastic pollution can alter natural processes, and habitats, reducing ecosystems’ ability to adapt to climate change, and food production capabilities, affecting millions of people’s livelihoods, and social well-being. A comprehensive study conducted across 60 major cities in India by the Central Pollution Control Board (CPCB) reveals that these urban centers collectively generate approximately 4059 metric tons per day (T/day) of plastic waste. Disparities exist in the proportion of plastic waste within the total Municipal Solid Waste (MSW), ranging from 3.10% in Chandigarh to 12.47% in Surat. On average, 6.92% of MSW is comprised of plastic waste. The detailed statistics pertaining to the generation of plastic waste in these 60 major Indian cities are appended herewith. Extrapolating from the data derived from these cities, it is estimated that India is poised to generate about 25,940 T/day of plastic waste. The ecological ramifications of plastic are noteworthy, given its propensity to contaminate air, water, and land due to the predominance of deleterious chemicals (Mageswari et al., 2018).

Utilizing plastic bricks represents a viable approach for the repurposing of plastics, thereby mitigating their wastage and mitigating associated pollution. Our focus centers on employing this methodology for plastic reuse in brick production. Notably, plastic bricks exhibit enhanced durability owing to the fibrous nature of plastic, and the incorporation of sand serves to prevent the formation of air pockets within the bricks. This results in a superior compression strength compared to concrete paving stones, which are prone to fracturing under heavy forces or prolonged exposure to adverse weather conditions.

The roadways within the Indian transportation system witness a pronounced increase in accidents due to inadequate lighting sources. This lack of proper illumination poses a significant risk to overall road safety, resulting in a heightened occurrence of vehicular collisions, especially in rural and underdeveloped areas. The absence of adequate lighting impairs visibility and complicates navigation for road users, making timely detection of potential hazards or obstacles challenging. Addressing these issues, the suggestion to incorporate the glowing plastic bricks emerges as a prudent solution to enhance nighttime visibility on highways. This involves strategically applying glowing plastic bricks on road dividers and speed breakers, thereby improving the visibility of these elements, such as dividers, turns, and speed breakers, during nighttime travel.

*Pseudomonas aeruginosa*, a well-known encapsulated, gram-negative, facultative aerobic, rod-shaped bacterium, is characterized by its pigmented nature. Various Pseudomonas species demonstrate the production of fluorescent yellow-green siderophores, such as pyoverdine (fluorescein), in response to iron-deficient environments (Marschner and Crowley, 1998). These pigments serve a dual purpose, being synthesized for iron acquisition and environmental protection against UV radiation (Burke et al., 1990). Harnessing the innate fluorescent property of *Pseudomonas aeruginosa*, we propose leveraging this biological mechanism as an eco-friendly alternative to radium paints. Radium paints, due to their high radioactivity, pose substantial health risks to workers and individuals handling radium-based products. Radium, mimicking calcium in the human body, tends to accumulate in the bones of exposed individuals, leading to the development of bone cancer, particularly in the jaws. In contrast to traditional glow-in-the-dark paints that rely on chemical formulations, our approach involves utilizing the natural and non-toxic fluorescent property of pyoverdine extracted from *Pseudomonas aeruginosa*. This biological strategy aims to imbue plastic bricks with a luminescent quality, providing a safer solution for both living organisms and the environment.

## Results and Discussion

### Assortment of plastic bricks

The cumulative volume of collected plastic waste amounted to approximately 5 kilograms, with an approximate requirement of 1 kilogram of plastic waste for the production of a single brick. The sourced plastic waste originated from diverse sources within our academic institution, Indrashil University, located in Kadi, Mehsana, Gujarat, encompassing the college campus, canteen, and laboratory facilities. Thermoplastics, predominantly employed in packaging and textile fibers, constitute approximately 80% of the overall plastic consumption. Nearly half of these thermoplastics are utilized in single-use applications. The remaining 20% to 25% of thermoplastics are directed towards long-term infrastructural purposes, such as pipes, cable coatings, and structural materials. The residual thermoplastics find application in durable consumer goods like furniture, automobiles, and other products characterized by an intermediate lifespan. Consequently, thermoplastic sets emerge as the most extensively utilized and readily available category of plastics. (Kazemi et al., 2021).

### Preparation of the mold and melting of plastic waste

The mold, constructed from iron material, possessed dimensions of 20 cm in length and 15 cm in breadth (Fig. 1a). The structural integrity of the mold was reinforced with a robust iron composition, ensuring its resilience to withstand the intense heat generated by the molten plastic material during the molding process. The gathered plastic waste underwent size reduction through cutting into smaller fragments. A container was set on high flame, as the container was hot enough around 180ºC, the pieces of plastic were added and flamed until it turned into a liquid state (Fig 1 b, 1c).

**Figure 1:**
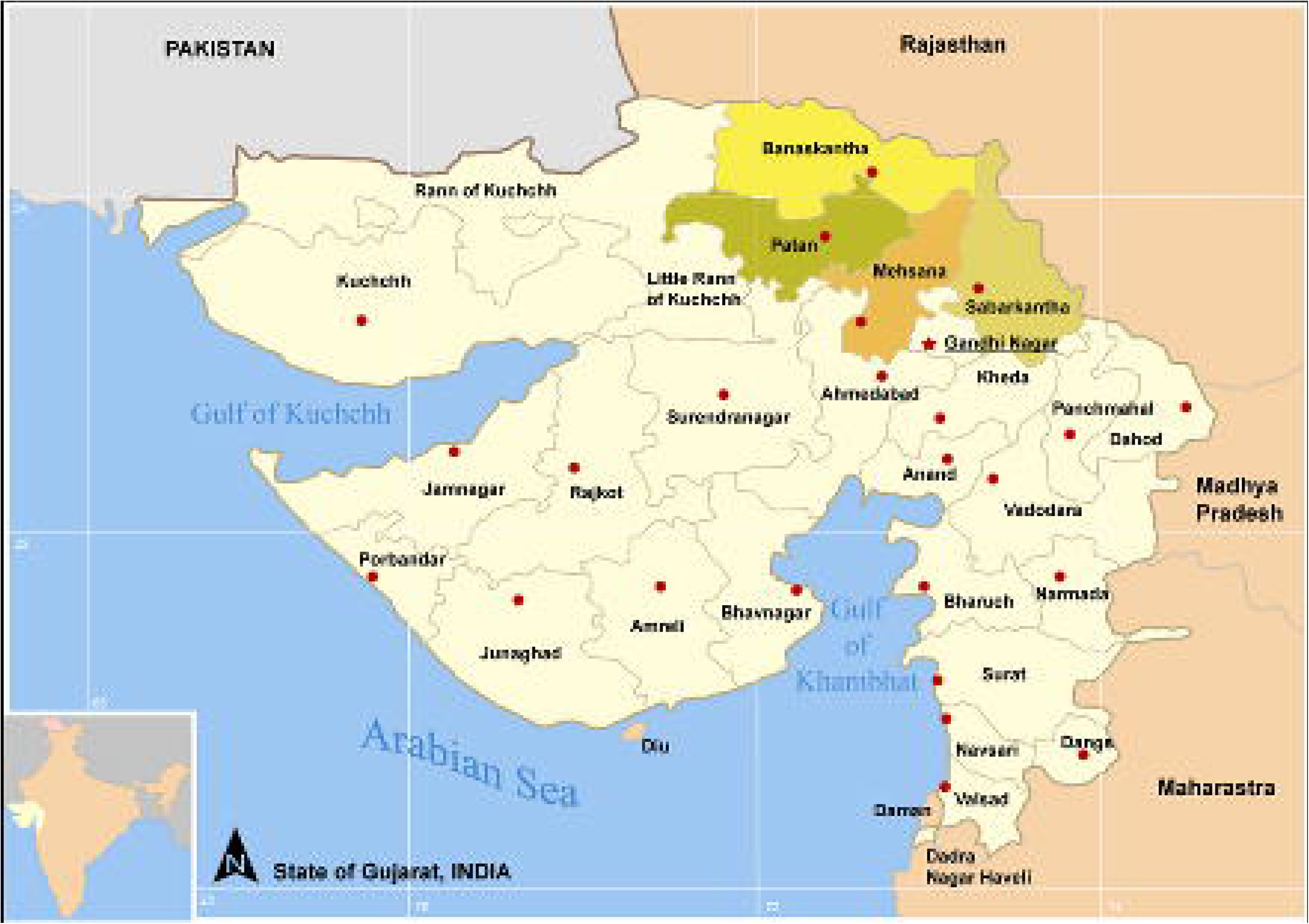
Steps of plastic brick preparation (1a) The prepared iron mold, (1b) Melting of plastic waste, (1c) Melted liquid form of the plastic waste, (1d) The basic structure of plastic brick.

### Molding of the plastic brick

The resulting plastic brick served as the foundational structure for the evaluation of our pigment; it demonstrated robustness against external forces and breakage (Fig. 1d). While the finishing did not match the standards observed in commercially available bricks due to the lab-based manufacturing approach, the plastic brick provided a platform for innovative applications in plastic utilization. The methodology proved to be convenient and executable, as noted by Uvarajan et al. (Uvarajan et al., 2022). This approach represents a unique integration of reuse and inventive design. Despite the handmade nature of the bricks using laboratory resources, they exhibited strength and resilience, remaining impervious to breakage. Waste reduction practices, encompassing reuse, recycling, processing, and storage, are widely recognized for their environmental, ethical, and financial merits across industries. To align with specified standards, it is crucial to assess the environmental friendliness, cost-effectiveness, and lightweight characteristics of construction materials. Plastic, with its diverse attributes such as durability, corrosion resistance, insulation properties for temperature and sound, energy efficiency, cost-effectiveness, long lifespan, and low weight (Haque, 2019), emerges as a versatile material for construction purposes. Hence, the introduction of the plastic brick concept represents a simplified yet impactful strategy in construction activities.

### Compressive strength

The capacity of a brick to withstand compression, as determined through testing on a compressive testing machine (CTM), is defined as compressive strength. Compressive strength is indicative of a material’s ability to resist failure in the form of cracks and fissures. The compressive strength of our two sample bricks was measured at 10.8 kN/min and 9.6 kN/min, respectively. Notably, the compressive strength of a material is influenced by its structural integrity and resistance to deformation under applied pressure. Our sample bricks exhibited lower compressive strength compared to standard plastic bricks, attributed to factors such as smaller brick size and the experimental nature of the trial conducted in a laboratory setting. During the test, the bricks are subjected to compression on both faces, and the maximum compressive load they can withstand before exhibiting cracks is observed and documented.

### Water absorption

Bricks undergo water absorption tests to evaluate their durability attributes, encompassing quality, firing intensity, and resistance to weathering. A brick’s susceptibility to damage from freezing is inversely correlated with its water absorption rate, with bricks exhibiting less than 7% absorption considered more resilient. This test assesses the compaction level of bricks, as their pores serve as conduits for water absorption. Increased pore density correlates with heightened water absorption. Bricks with absorption rates below 3% are categorized as vitrified. In our study, sample bricks exhibited water absorption percentages of 0.21% and 0.13%, respectively. During the test, bricks are initially weighed in a dry state, then submerged in fresh water for a 24-hour period. Subsequently, the bricks are removed from the water, dried to remove excess moisture, and weighed in their wet state. The discrepancy in weight indicates the amount of water absorbed by the brick. Superior-grade bricks exhibit minimal water absorption, with high-quality specimens absorbing no more than 20% of their weight.

### Selection of bacterial strain and culture medium

The *Pseudomonas aeruginosa* strain was selected for the production of the desired pyoverdine pigment (Fig. 2a & 2b). Various strains of Pseudomonas are also available for pyoverdine pigment production. For instance, the yellow-green, fluorescent, water-soluble pigment from *Pseudomonas fluorescens* was initially characterized as “pyoverdine” by Turfreijer in 1941 (Turfreijer 1941). This terminology was chosen in analogy to the phenazine pigment pyocyanine, which is synthesized by *Pseudomonas aeruginosa*. Alternative terms such as “bacterial fluorescein” and “fluorescin” have been used occasionally in literature, as preferred by King et al. and Lenhoff (King et al., 1948; Lenhoff, 1963). Pyoverdine has been further expanded by other authors like Elliot and Hulcher to encompass any pigment produced by luminous pseudomonads (Elliot, 1958; Hulcher, 1968). However, it may be more desirable to designate pigments of this class with a suffix denoting the species that produces them, such as pyoverdinePf for the pyoverdine produced by *Pseudomonas fluorescens*, given that prior unpublished studies have indicated structural differences among pigments from various species of fluorescent pseudomonads. (Wu et al., 2015) (Pratt, 1993) (Curtiss, 1942) (Ganne et al., 2017) (Burke et al., 1990)

**Figure 2:**
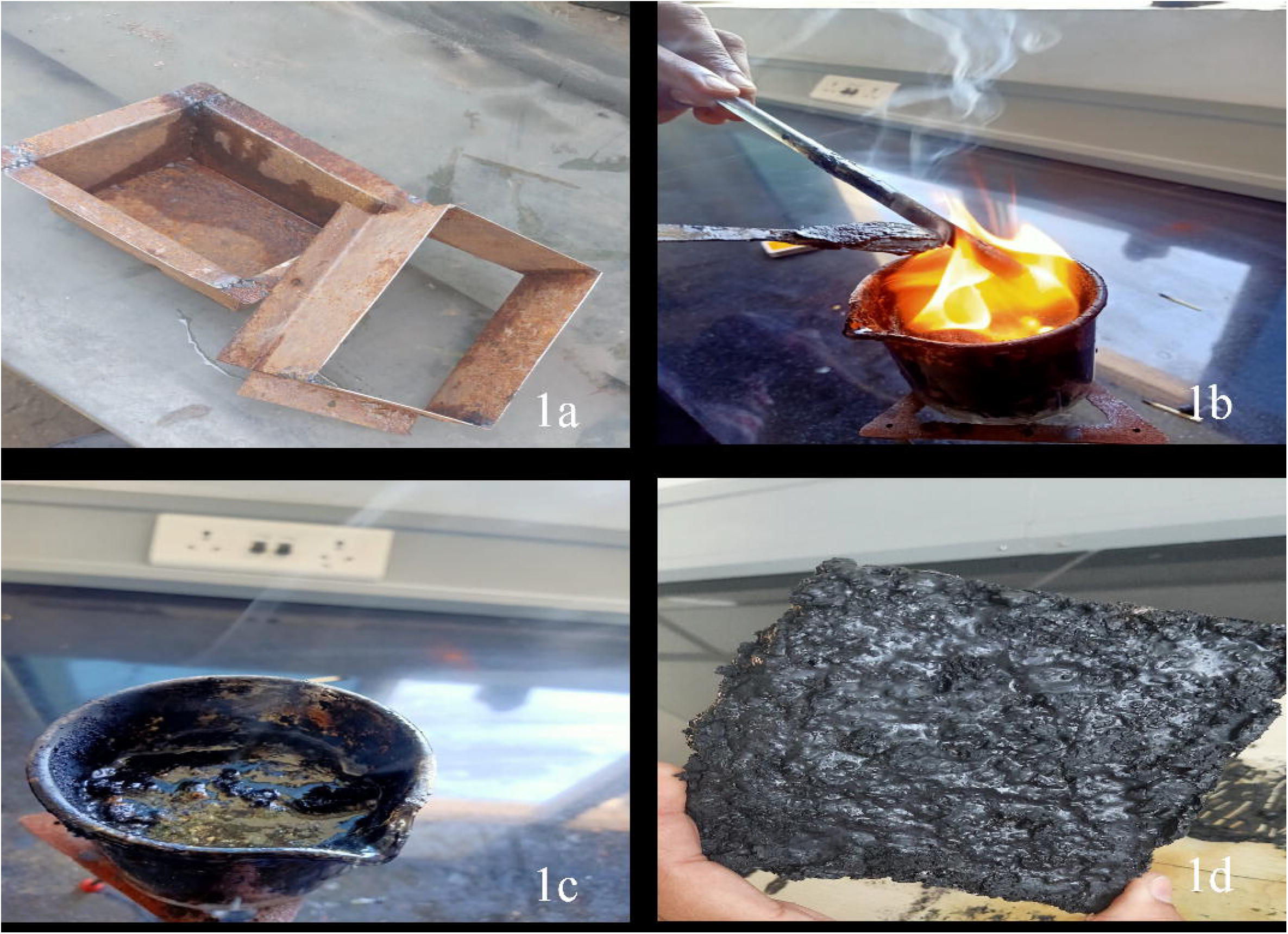
Extraction of pyoverdine pigment, (2a) Isolated *Pseudomonas aeruginosa* colonies on kings’ media agar plates, (2b) Kings Broth inoculated with selected *Pseudomonas aeruginosa* strain, (2c) Supernatant saturated with 0.5g of NaCl and half the volume of CHCl3/ Phenol (1:1), (2d) Extracted Pyoverdine pigment under UV light.

### Extraction of Pyoverdine pigment

The extraction of Pyoverdine pigment was executed with a systematic approach (Fig. 2c & 2d). The fluorescent characteristics of this siderophore prompted consideration for its potential as a replacement for radium paints (Wu et al., 2015), owing to its luminescent properties. Moved by this attribute, we endeavored to extract and utilize the pigment as a glowing substitute, aiming to mitigate the hazards associated with radium paints. Pseudomonas aeruginosa pigment was selected for our investigation due to its inherent fluorescence, presenting a safer and more natural alternative to hazardous chemicals. Radium possesses numerous hazardous properties; it is a component of self-luminous paints utilized for dials, which may lead to “radium poisoning” in individuals exposed to such paints. Radium emits three types of radiation: alpha, beta, and gamma rays (Pratt, 1993). Alpha rays, consisting of doubly charged helium atoms, have limited penetrating power and primarily affect the blood-forming marrow upon ingestion. Beta rays, high-speed electrons with greater penetrating capabilities, can cause skin burns and exert effects from outside the body. Gamma rays, akin to X-rays, penetrate deeply into tissues, inducing slow changes in blood structure detectable through blood counts. Radium decay generates radon gas, which rapidly dissipates upon inhalation. However, once ingested, radium remains in the body, manifesting its deleterious effects over time (Curtiss, 1942). Our exploration into the adverse effects of radium prompted the conception of glowing plastic bricks utilizing the fluorescent pigment of *Pseudomonas aeruginosa*. Siderophores, produced by *Pseudomonas aeruginosa* under iron-deficient conditions (Ganne et al., 2017), not only serve as iron scavengers but also confer protection against UV radiation (Burke et al., 1990).

### The glowing plastic bricks

The extracted pyoverdine pigment was applied to the bricks via spraying, followed by exposure to sunlight for drying to facilitate UV light absorption from solar radiation (Fig. 3a). Subsequently, the bricks were evaluated under flashlights in darkness, revealing a fluorescence phenomenon (Fig. 3b). These luminescent plastic bricks can be utilized for road striping or road markings, which are vital components of traffic management and road safety.

**Figure 3:**
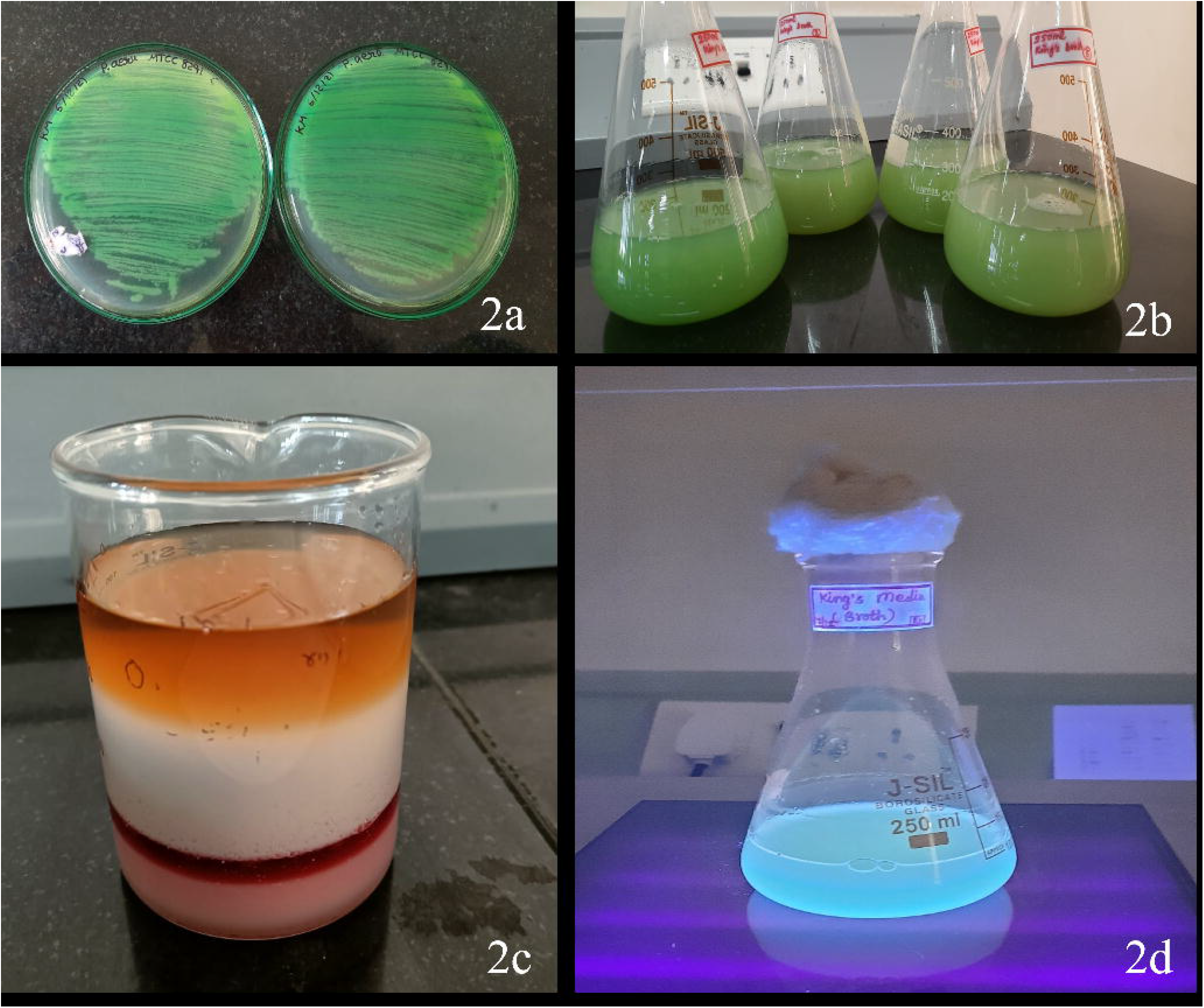
Glowing plastic bricks(3a) Pyoverdine pigmented plastic brick (under UV light) and (3b) Pigmented brick under street light.

## Conclusion

The study aimed to achieve luminescence in plastic bricks, illustrating the reuse of waste materials. Considering the pervasive use of plastics and their persistent non-biodegradability, repurposing them into plastic bricks is a crucial strategy to mitigate plastic pollution and foster sustainable development. Additionally, our objective was to impart luminescence to these bricks using biological methods, specifically by extracting fluorescent pigment from *Pseudomonas aeruginosa*, thereby circumventing the use of chemical alternatives. The extraction process from *Pseudomonas aeruginosa* was intricate yet successful, with the obtained pigment subsequently applied to the plastic brick via a spraying technique. Our aim was to obviate the necessity for hazardous chemicals commonly employed to induce fluorescence, such as Radium, glow-in-the-dark materials, and Radium-based paints and sprays, due to their carcinogenic properties. Consequently, we endeavored to introduce an alternative approach to fluorescence. These glowing plastic bricks can be utilized for road striping or road markings, which are crucial elements of traffic management and road safety.

## Materials and Methods

### Assortment of plastic waste

Our objective was to devise a straightforward methodology for the production of plastic bricks. In the initial phase, we commenced the acquisition of waste plastics sourced from diverse locations within our academic institution (Indrashil University) situated in Kadi, Mehsana, Gujarat, including the college campus, canteen, and laboratories (Fig. 4). The aggregate weight of the collected plastic waste approximated 5 kilograms, with the requisite and utilized quantity of plastic waste for the creation of one brick being approximately 1 kilogram. Subsequently, we amalgamated the collected plastics and categorized them based on their composition as either Thermoplastics or Thermosetting plastics. Rigorous washing procedures were employed on all the accumulated and designated plastic waste to ensure the elimination of contaminants such as dirt and other impurities.

**Figure 4:**
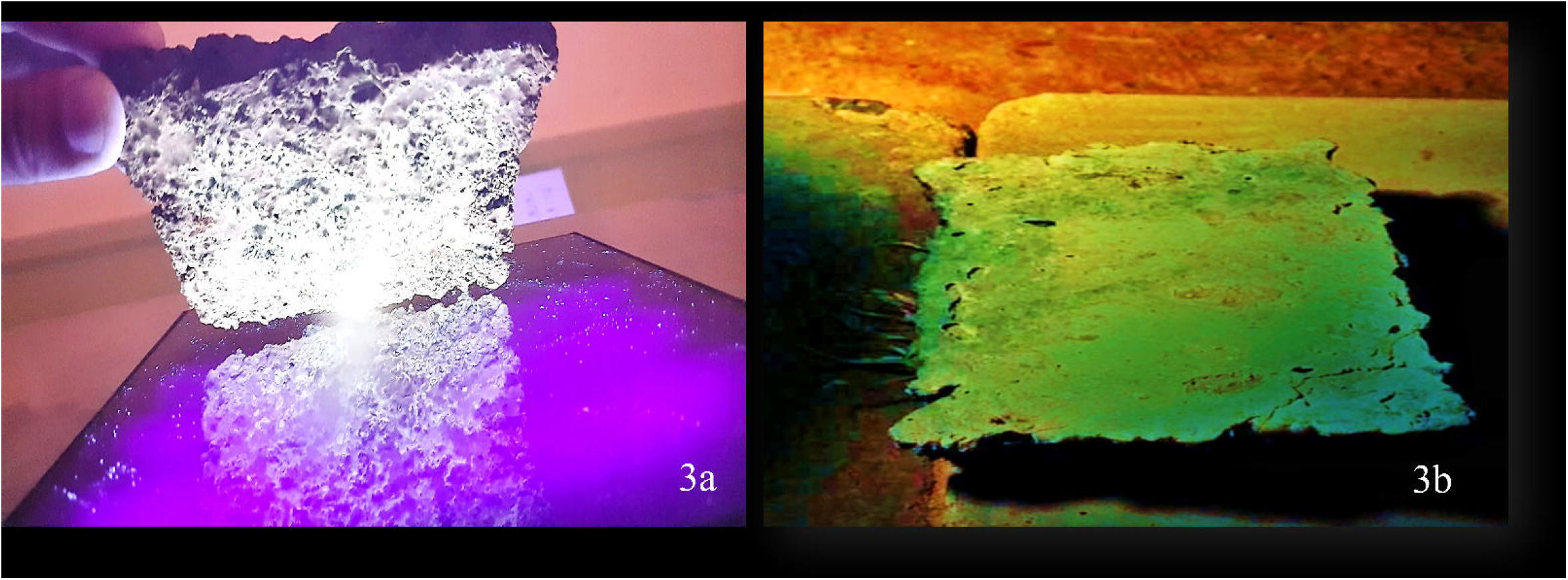
Map of Gujarat highlighting Mehsana district

### Preparation of the mold

The mold employed in this study was constructed from iron material, featuring dimensions of 20cm in length and 15cm in breadth. To ensure structural integrity and resistance to the elevated temperatures associated with the molten plastic, the mold was fortified with a substantial iron load. This reinforcement was essential to withstand the thermal stress induced by the molten plastic during the molding process.

### Melting of plastic waste

The collected plastic waste underwent initial processing by being reduced into smaller fragments. Subsequently, a receptacle was exposed to a high-temperature flame, reaching approximately 180ºC. Once the container attained the required temperature, the aforementioned plastic fragments were introduced and subjected to the heat until they underwent a phase transition, transforming into a molten state.

### Molding of the plastic brick

The molten thermoplastic material was introduced into the designated mold and allowed to undergo solidification over approximately 24 hours. The liquid state, derived from the pyrolysis of plastic waste, necessitated immediate pouring at an elevated temperature to prevent premature solidification, ensuring adherence to the container walls. Following the 24-hour solidification period, the resultant bricks were demolded, measuring 17 cm in length and 15 cm in breadth. Subsequent refinement procedures were executed to ameliorate surface irregularities, resulting in final brick dimensions of 15 cm in length and 10 cm in breadth. Post-finishing, the bricks were prepared for the application of fluorescent pigment named pyoverdine.

### Compressive strength

The computation of the compressive strength of the specimen brick was determined by the application of the following formula:

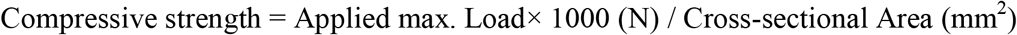

Specimen bricks are subjected to compressive strength tests using a compression testing machine with a capacity of 3000 kN (Kilo Newtons). The bricks undergo uniform loading at a rate of 2.9 kN/min. Upon reaching the maximum load, it is inferred that the applied load has caused no further escalation in the indicator reading on the testing equipment, signifying failure of the specimen under test conditions.

### Water absorption

The water absorption capacity of bricks should not surpass 12% of their total weight. To evaluate water absorption, test bricks undergo a standardized procedure. Initially, the bricks are desiccated in an oven at temperatures ranging from 105°C to 115°C until a consistent weight is attained. Subsequent to desiccation, the bricks are allowed to cool to ambient temperature and are weighed (W1). These thoroughly desiccated bricks are then submerged in clean water for a period of 24 hours, maintaining a temperature between 20°C and 27°C. Upon completion of the immersion period, the bricks are withdrawn, any residual water is removed, and they are promptly reweighed (W2).

The computation for water absorption, presented as a percentage relative to weight, is determined by the formula:

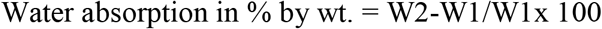

### Culture of *Pseudomonas aeruginosa*

The *Pseudomonas aeruginosa* strain was acquired from Microbial Type Culture Collection and Gene Bank, Chandigarh, India. The preparation of nutrient broth, nutrient agar plates, and glycerol stock solution ensued, followed by autoclaving. The lyophilized culture was meticulously opened according to the prescribed protocol and activated using nutrient broth. Subsequently, the *Pseudomonas aeruginosa* culture was streaked onto nutrient agar plates, which were then incubated at 25°C for 24 hours. After the incubation period, growth was assessed in the broth through turbidity measurements and observation of colonies on the agar plates.

The broth, containing viable culture, was cryopreserved using glycerol stock solutions at -80°C, while the agar plates with colonies were stored under refrigerated conditions. Kings’ media was formulated and utilized for streaking isolated colonies from the nutrient agar plates, followed by another 24-hour incubation at 25°C. Subsequently, the plates were refrigerated. To initiate pigment production, 100 ml Kings Broth was prepared for inoculation with colonies obtained from the nutrient agar plates. Following autoclaving, the broth was cooled to room temperature, and the colony was inoculated and subjected to shaking conditions at 25°C.

Upon completion of the 24-hour incubation, the optical density was measured at 600nm, yielding an absorbance value of 1.03. To facilitate further inoculation, 5 flasks of 250 ml Kings Media were prepared with a 5% culture concentration (v/v). This concentration corresponds to 12.5 ml of the previously cultured 100 ml in each flask, as outlined by King et al. (King et al., 1948).

### Extraction protocol of pyoverdine pigment

A quantity of 50 mg (0.05 g) of FeCl3 was introduced to the culture, initiating centrifugation in accordance with the methodology outlined by Waring et al. (Waring et al., 1942). Subsequently, the pellets were discarded, and the supernatant was retrieved. The Supernatant was subjected to saturation with 0.5 g of NaCl, and half the volume of CHCl3/Phenol (1:1) was incorporated. Extraction was performed using a sonicator for a duration of 3 minutes, comprising 1-minute intervals repeated three times with 5 minutes of rest in between. The resulting organic phase was collected, followed by the addition of a double volume of diethyl ether and half the volume of distilled water. The sample underwent evaporation in a rotary evaporator, with absorbance measurements taken at 400 nm as per the methodology established by Meyer et al. (Meyer et al., 1978).

A similar extraction process for pigment was employed for all five flasks on their respective days. The extracted pigments were shielded with aluminum foil and left for natural evaporation of any residual water content. The pigment was subsequently applied to the plastic brick and exposed to an open area, facilitating drying and absorption of ample UV light from the atmosphere.

### Coating of pyoverdine pigment on the plastic bricks

The pyoverdine pigment was collected, and the initial layer was applied via spraying onto the plastic bricks. Following the pigment application, the bricks were left undisturbed to facilitate absorption, subsequently placed in an open area exposed to natural sunlight, such as an open terrace, ensuring ample sunlight exposure and out of reach from children or any animal groups.

After the complete drying and absorption of the first layer of pigment onto the brick, a second layer was sprayed onto the surface. Once again, the bricks were positioned under sunlight to enable drying.

## Acknowledgement

We acknowledge student startups and innovation policy, Government of Gujarat for providing financial support to carry out this study.

